# Genomic insights on the potential role of the accessory genome in the emergence of a novel geographically restricted *K. pneumoniae* lineage as a high-risk clone

**DOI:** 10.1101/2024.03.18.585244

**Authors:** Nathália Bighi, Fernanda Freitas, Sérgio Morgado, Rosângela Cipriano, Ana Carolina Vicente, Érica L. Fonseca

## Abstract

*Klebsiella pneumoniae* causes life-threatening nosocomial infections and is featured by a remarkable propensity for multidrug resistance acquisition. Infections caused by multidrug- (MDR) and extensively drug-resistant (XDR) strains lead to a limitation of therapeutic options and an increase in persistent infections, and they are usually represented by high-risk lineages. Based on these features and their relevance to global public health, most of the studies focused on such high-risk clones, and little is known about the epidemiological and evolutionary dynamics of new/geographically restricted lineages. This study aimed to unveil the antimicrobial resistance and virulence genetic repertoire of a clinical XDR *K. pneumoniae* (Kp199) strain belonging to geographically-restricted ST, not linked to any known clonal complex. Its intrinsic (*gyrA, parC, ramR, soxR* and *soxS* mutations) and acquired resistome agreed with the observed XDR phenotype. An extensive arsenal of both antibiotic and heavy metal resistance genes was observed, as well as genes involved with resistance to several antiseptics currently used in clinical settings. The co-occurrence of *bla*_KPC-2_ and *bla*_NDM-1_ carbapenemase genes in Kp199 was an alarming finding since it could contribute to increased carbapenem resistance. Kp199 virulome was associated with bacterial survival and replication during infections. This study raises concern about a novel, geographically restricted *K. pneumoniae* lineage harbouring a huge resistome and virulome, which may strongly contribute to its successful establishment as an epidemic lineage. Therefore, our findings underscore the importance of vigilant surveillance and control measures to mitigate the threat posed by the potential emergence of new high-risk pandemic clones.

## INTRODUCTION

*Klebsiella pneumoniae* is a prominent opportunistic pathogen considered one of the most challenging agents of nosocomial infections, which are mainly caused by multidrug resistant *K. pneumoniae* (MDR-KP). The evolution of this multidrug resistance in association with the acquisition of virulence factors by horizontal gene transfer contributes to its persistence in clinical settings and to the efficient establishment of infections, leading to the emergence of high-risk pandemic clones [1]. In fact, most reports on MDR-KP isolates include successful lineages, such as ST15, ST512, and those belonging to clonal group CG258, ST11 and ST258, which are currently the major lineages responsible for the global dissemination of carbapenemase-coding genes *bla*_KPC-2_ and *bla*_NDM-1_ and carbapenem resistance [2]. Such carbapenem-resistant high-risk clones often combine resistance to last-resort antibiotics, including tigecycline and polymyxins, leading to the emergence of extensively drug resistant (XDR) or pandrug resistant (PDR) strains [3]. However, there is a bias regarding the identification and characterization of the genetic components involved with the resistance and dissemination of pandemic clones, since most reports focus on known high-risk pandemic lineages due to their impact on global public health. In this context, little is known about the genetic arsenal involved with antibiotic resistance, virulence and persistence potential of newly emergent or geographically restricted *K. pneumoniae* lineages with a high propensity of spreading. This scarcity of information hinders the understanding of the emergence of high-risk strains in the future, demonstrating the need for epidemiological surveillance of such lineages with local coverage. Therefore, this study performed a comprehensive genomic analysis to better understand the pandemic potential of a newly emerging, extensively resistant *K. pneumoniae* lineage (Kp199) recovered in Amazon region co-producing *bla*_NDM-1_ and *bla*_KPC-2_ carbapenemase genes.

## MATERIAL AND METHODS

The Kp199 strain was recovered in 2022 from a blood infection case in a clinical setting in the Amazon Region (Maranhão). The antimicrobial susceptibility test to all antibiotics considered for *K. pneumoniae* resistance classification [4] was determined by the disk-diffusion method: ampicillin/sulbactam, amoxicillin/clavulanate, piperacillin/tazobactam, cefazoline, cefotetan, cefoxitin, cefuroxime ceftazidime, ceftriaxone, cefepime, ceftaroline, aztreonam, meropenem, imipenem, ertapenem, doripenem, netilmicin, amikacin, gentamicin, tobramycin, tetracycline, minocycline, doxycycline, tigecycline, trimethoprim/sulfamethoxazole, chloramphenicol, fosfomycin, ciprofloxacin, and colistin.

The results were interpreted according to the Clinical and Laboratory Standards Institute (CLSI) [5], and European Committee on Antimicrobial Susceptibility Testing (EUCAST) (for tigecycline and colistin) guidelines [6]. The Colistin MIC was determined by broth microdilution (MIC breakpoint for resistance >2 μg/mL).

The Kp199 genome was obtained on the Illumina Hiseq 2500 using Nextera paired-end library kit for library construction. SPAdes assembler v3.15.2 (https://github.com/ablab/spades/) was used for genome assembling. Gene prediction/annotation was conducted with Prokka v1.14.6 (https://github.com/tseemann/prokka). Core genome MLST (cgMLST) was determined in the Bacterial Isolate Genome Sequence Database (BIGSdb; http://bigsdb.pasteur.fr/klebsiella/). It was also submitted to PathogenWatch, a global platform for genomic surveillance (https://pathogen.watch/).

To compare Kp199 with available genomes of *K. pneumoniae* worldwide, a total of 60,437 *K. pneumoniae* genomes were retrieved from GenBank. The Mash v2.2.2 software (https://bioweb.pasteur.fr/packages/pack@Mash@2.2) was used to estimate the distance between the Kp199 genome and those available in the database and identify the closest genomes for subsequent phylogenomic analysis. The core genome was determined using Roary v3.13.0 (https://github.com/sanger-pathogens/Roary), and the single nucleotide polymorphisms (SNPs) sites of the concatenated core genes were extracted using SNP-sites v2.5.1 (http://sanger-pathogens.github.io/snp-sites/). A maximum likelihood tree with 1,000 ultrafast bootstrap replicates, employing the substitution model VMe+ASC+R3, was generated by IQTree v1.12 and visualized using iTOL (https://itol.embl.de/).

Plasmid replicons, integrative and conjugative elements (ICEs), antimicrobial resistance determinants and virulence factors were identified using PlasmidFinder v.2.0 (https://cge.cbs.dtu.dk/services/PlasmidFinder/), ICEfinder (https://bioinfo-mml.sjtu.edu.cn/ICEfinder/index.php), the Comprehensive Antibiotic Resistance Database (CARD - https://card.mcmaster.ca/), and Virulence Factors Database (VFDB - http://www.mgc.ac.cn/VFs/), respectively. The Kaptive tool was used for polysaccharide locus typing of bacterial capsules (https://kaptive-web.erc.monash.edu/).

The deduced protein of each Kp199 chromosomal gene involved with fluoroquinolones (*gyrA* and *parC*), and tigecycline resistance (*ramR, ramA, soxR, soxS, marA, marR*, and *acrR*) was compared with that of the wild-type reference strain *K. pneumoniae* NTUH-K2044 (NC_012731.1).

## RESULTS AND DISCUSSION

The *in vitro* analyses demonstrated that Kp199 corresponded to an extremely drug-resistant (XDR) strain, being resistant to all tested antibiotics, except for tetracycline and colistin. The MLST revealed that Kp199 belonged to ST3128, a geographically restricted lineage, only reported until now in a broiler production chain in Germany between 2014-2016 [7]. Interestingly, although belonging to ST3128, the phylogenetic reconstruction performed with Kp199 genome and the closest publicly available *K. pneumoniae* genomes revealed that Kp199 was genomically distinct from ST3128 and unrelated to known clonal complexes (Figure 1), demonstrating that it corresponded to a new lineage.

**Figure 1:**
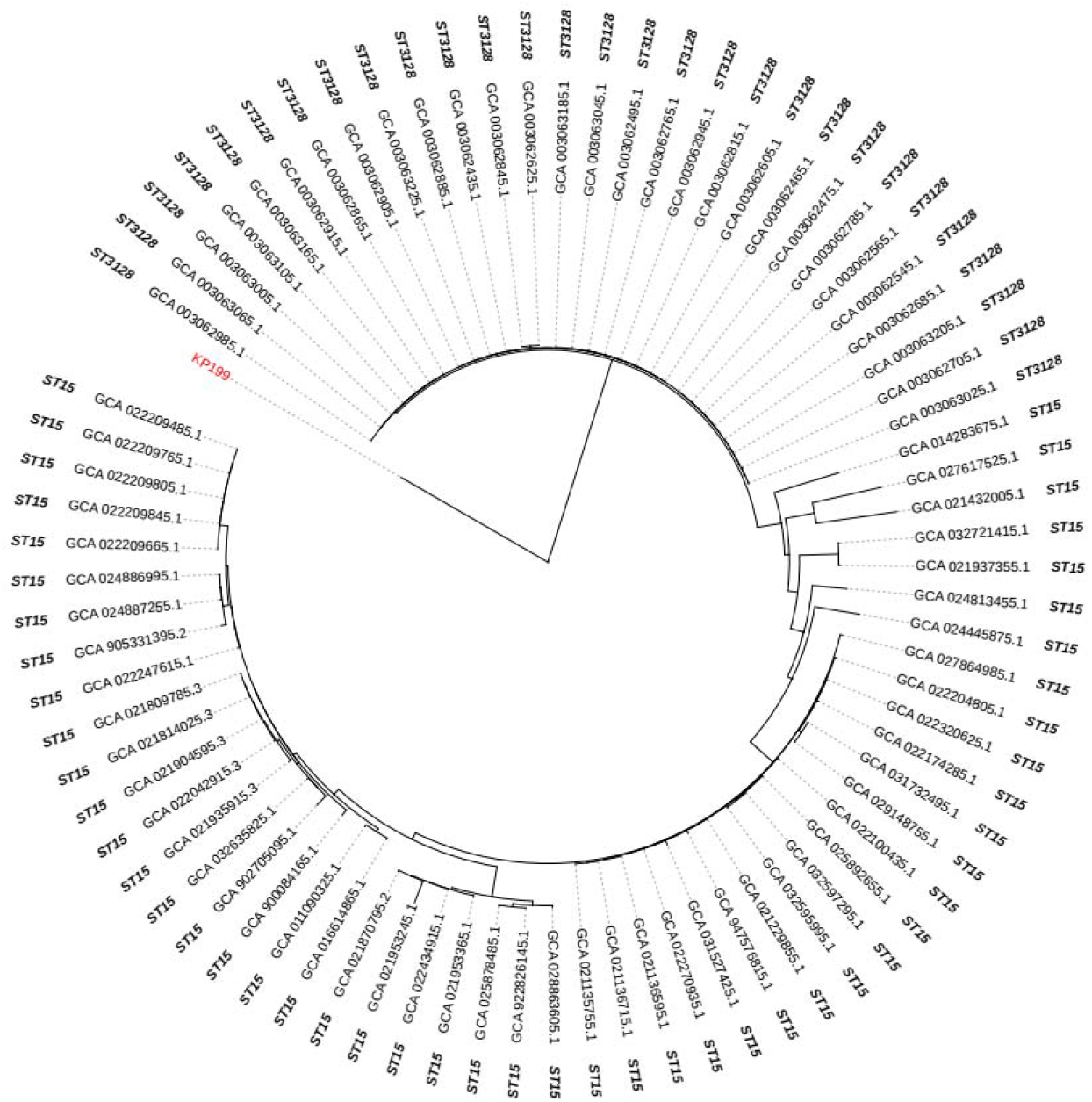
Maximum likelihood tree based on the core genome of KP199 (red) and the closest *K. pneumoniae* genomes available in Genbank. The corresponding sequence types (STs) are indicated.

The Kp199 resistome mining revealed the presence of a remarkable arsenal of antimicrobial resistance genes (ARGs) and modifications in housekeeping genes, explaining the observed XDR phenotype. The intrinsic Kp199 resistome related to fluoroquinolones (FQ) revealed the presence of substitutions in the quinolone resistance-determining regions (QRDR) of GyrA (S83F and D87A) and ParC (S80I) already known to be involved with FQ resistance. Considering tigecycline resistance, although this phenotype is still rare in *K. pneumoniae, acrAB* efflux pump overexpression in response to alterations in their regulatory genes (*ramR, ramA, soxR, soxS, marA, marR, acrR, and rarA*) is the most common mechanism responsible for this resistance phenotype [8]. Among the aforementioned genes involved with tigecycline resistance, only *ramR, soxR* and *soxS* were altered. The Kp199 RamR deduced protein presented the substitutions A19V, which has no significant effect on tigecycline susceptibility or the upregulation of *ramA* and *acrA* [9], and K63M, which, to our knowledge, has never been reported before. The SoxR presented four novel substitutions (C124W, D137G, K138D, E142K), not associated with resistance yet. However, a previous study demonstrated that the substitution of a glutamic acid (E) at position 142 could be involved with *soxS* overexpression and consequently, with tigecycline resistance emergence [10]. Similarly, SoxS also harboured a new amino acid substitution (Q70L) never reported elsewhere.

Kp199 acquired resistome was composed of ARGs associated with resistance to β-lactams (*bla*_SHV-11_, *bla*_CTX-M-15_, *bla*_OXA-9_, *bla*_OXA-1_, *bla*_TEM-1_), aminoglycosides (*aacA4, aadA1, aph(3’)-VI, aac(3)-IId*), streptothricin (*sat1*), bleomycin (*ble*) fluoroquinolones (*qnrS1, qnrE1, oqxAB*), trimethoprim (*dfrA15*), Quaternary ammonium compounds (*qacE*Δ*1*), sulfonamides (*sul1, sul2*), rifampin (*arr-3*), chloramphenicol (*catB3, cmlA4*) and carbapenems (*bla*_NDM-1_, *bla*_KPC-2_). It is worth mentioning that the co-occurrence of NDM-1 and KPC-2 is not a common finding since only a few studies reported the simultaneous production of these two carbapenemase genes in *K. pneumoniae*. Interestingly, such reports were restricted to pandemic clones occurring in Asia, mainly in China mostly represented by the hypervirulent/multidrug resistant ST11-K64 [2]. Curiously, outside China, the only two reports of *K. pneumoniae* co-harbouring *bla*_NDM-1_/*bla*_KPC-2_ were in the South America: in a pandemic ST258 strain in Uruguay [11] and in a new ST in Brazil, as found here for Kp199 [12]. The co-existence of NDM-1 and KPC-2 is a worrisome association since it accounts for increased carbapenem MICs due to the synergistic effect of both enzymes as observed previously [2]. Moreover, both carbapenemase genes were found in the context of mobile elements in Kp199. The *bla*_NDM-1_ was present in a genomic segment of 13 kb that also harboured the ARGs *qnrS1, aph(3’)-VI* and *ble* and shared 100% nucleotide identity and gene synteny with several plasmids of *K. pneumoniae* and other *Enterobacteriaceae* worldwide.

The *bla*_NDM-1_ was immediately downstream of IS*Aba125* as found previously, but in this case, this element was disrupted by IS*630*. Similarly, the *bla*_KPC-2_ gene was flanked by IS*6*-like elements. In fact, except for *bla*_SHV-11_, which is constitutively encoded in *K. pneumoniae* species, all other predicted ARGs were also found in the context of distinct mobile platforms, such as insertion sequences and plasmid-related segments carrying the plasmid replication initiation *rep* genes characterizing IncFIIK, IncFIB, IncFII and IncHI1B, increasing the chances of ARG horizontal transmission. For example, the ARGs *bla*_OXA-9_, *qacE*Δ*1* and *sul1* were found embedded in a region of genomic plasticity of 31.4 kb also carrying several genes required for plasmid conjugal transfer (*mobHI* relaxases, and *tra* operon).

Disinfection by biocides is essential to control infections and the emergence of disinfectant-resistant microbes, mainly those already resistant to antibiotics, increases the risk of bacterial transmission in healthcare settings. Moreover, there is a relationship between biocide use and the emergence of both biocide and antibiotic resistance, particularly multidrug-resistant pathogens [13]. Similarly, the frequent use of heavy metal compounds in nature, such as in livestock, not only promotes the emergence of heavy metal resistance but also triggers the co-selection of antibiotic resistant bacteria [14]. In fact, besides ARGs, Kp199 also harboured genes involved with resistance to biocides and antiseptics frequently used in clinical settings, such as *cepA, fabI, kpnEF*, and *kmrA*, and an expressive set of heavy metal resistance determinants. This strain harboured genes associated with resistance emergence to nickel (*nikABCDER* operon and *rcnA* gene), arsenic (*arsABCDHR* and *pstSCAB*), copper/silver *(silESRCFBAP, cusRSCFBA, copGEABCDR, cpxPRA*), cobalt/cadmium/zinc (*czc* genes), mercury (*merRTPCADE*), magnesium/nickel/copper/cobalt (*corABCD*), zinc (*znuACB*), molybdenum (*modABCE*), copper/zinc (*mdtABC*), and tellurium (*terDABCDF*). Curiously, the presence of copper could not only co-select ARGs but also positively correlate with the acquisition of some virulence factors [15]. Interestingly, the *sil, cop*, and *ter* resistance operons found in Kp199 shared high identity (99%) and complete synteny with part of the 220 kb virulence plasmid pLVPK (GenBank accession no.

AY378100), previously identified in a bacteremic *K. pneumoniae* strain in Taiwan. This plasmid is known for harbouring the main genetic markers that characterize hypervirulent (presence of salmochelin *iroBCDEN*, aerobactin *iucABCDiutA*, yersiniabactin-coding *irp* and *ybt* genes, and colibactin *clbAR*) and hypermucoviscous (*magA, rmpA, rmpA2*) *K. pneumoniae* strains [16]. However, none of them were identified in Kp199.

Although Kp199 was not characterized as hypervirulent, it harboured several genes involved with iron uptake (*entHABECF, fepABCDG, entS* enterobactin siderophore-related genes), adherence (*ilpA*, Type 1 *fimBEAICDFGHK* fimbriae, and *ecpRABCDE* pilus biosynthesis operons), biofilm formation (*luxS*), lipid A biosynthesis (*lpxC*), and resistance to intracellular killing (*mgtC*). Other important virulence factors of *K. pneumoniae* are the capsular polysaccharide (CPS) K and the lipopolysaccharide (LPS) O antigens, which protect against phagocytosis, humoral defenses, and contribute to enhanced virulence in general [16]. The Kp199 has the K21/O1 serotype, which has been associated with outbreaks of respiratory and urinary nosocomial infections, but is not related to the capsule of hypervirulent strains, which usually belong to K1, K2, K28, K47, and K63 serotypes, although the O1 antigen has been previously described as a major contributor to the virulence of pyogenic liver abscess causing *K. pneumoniae* [17]. Therefore, despite not being a hypervirulent strain, Kp199 carries an important virulence arsenal that contributes to bacterial survival and multiplication during infections.

## CONCLUSIONS

The presence of a remarkable virulence and antibiotic resistance genetic repertoire in a novel lineage (Kp199), comparable with that of the pandemic clones such as ST258, highlights its potential success as a persistent nosocomial pathogen and raises the worrisome perspective of a lineage with local coverage to emerge as a global, extensively drug resistant clone in the future. Moreover, this study provides insights into the co-transmission risk of both *bla*_NDM-1_ and *bla*_KPC-2_ genes, contributing to the selection and worldwide dissemination of highly carbapenem-resistant strains that may severely compromise the treatment and outcome of the associated infections. Therefore, our findings strengthen the valuable contribution of continuous surveillance of extensively drug-resistant strains, even if they do not belong to pandemic lineages, to recognize the epidemiological profile of pathogens in clinical settings and avoid unfavorable clinical outcomes. Finally, the tight association of the virulence and antibiotic resistance determinants with regions of horizontal transfer stresses, one more time, that the clonal success of high-risk lineages is primarily linked to the acquisition and spread of mobile genetic elements.

## References

1. Di Pilato V, Pollini S, Miriagou V, Rossolini GM, D’Andrea MM (2024) Carbapenem-resistant Klebsiella pneumoniae: the role of plasmids in emergence, dissemination and evolution of a major clinical challenge. Expert Rev Anti Infect Ther. 10.1080/14787210.2024.2305854.

2. Sun S, Cai M, Wang Q, Wang S, Zhang L, Wang H (2023) Emergency of the plasmid co-carrying bla(KPC-2) and bla(NDM-1) genes in carbapenem-resistant hypervirulent Klebsiella pneumoniae. J Glob Antimicrob Resist. 36:26–32. doi: 10.1016/j.jgar.2023.11.008.

3. Lázaro-Perona F, Sotillo A, Troyano-Hernáez P, Gómez-Gil R, de la Vega-Bueno Á, Mingorance J (2018) Genomic path to pandrug resistance in a clinical isolate of Klebsiella pneumoniae. Int J Antimicrob Agents 52:713–718. doi: 10.1016/j.ijantimicag.2018.08.012.

4. Magiorakos AP, Srinivasan A, Carey RB, Carmeli Y, Falagas ME, Giske CG et al (2012) Multidrug-resistant, extensively drug-resistant and pandrug-resistant bacteria: an international expert proposal for interim standard definitions for acquired resistance. Clin Microbiol Infect 18:268–81. doi: 10.1111/j.1469-0691.2011.03570.x.

5. Clinical and Laboratory Standards Institute (2021) Performance standards for antimicrobial susceptibility testing. 32nd ed. CLSI supplement M100. Wayne, PA.

6. European Committee on Antimicrobial Susceptibility Testing (EUCAST) Breakpoint tables for interpretation of MICs and zone diameters (Version 11.0). 2021; https://www.eucast.org/ [accessed 12 December 2023].

7. Daehre K, Projahn M, Friese A, Semmler T, Guenther S, Roesler UH (2018) ESBL-Producing Klebsiella pneumoniae in the Broiler Production Chain and the First Description of ST3128. Front Microbiol 9:2302. 10.3389/fmicb.2018.02302.

8. Park Y, Choi Q, Kwon GC, Koo SH (2020) Molecular epidemiology and mechanisms of tigecycline resistance in carbapenem-resistant Klebsiella pneumoniae isolates. J Clin Lab Anal 34:e23506. doi: 10.1002/jcla.23506.

9. Chiu SK, Huang LY, Chen H, Tsai YK, Liou CH, Lin JC et al (2017) Roles of ramR and tet(A) Mutations in Conferring Tigecycline Resistance in Carbapenem-Resistant Klebsiella pneumoniae Clinical Isolates. Antimicrob Agents Chemother 61:e00391–17. doi: 10.1128/AAC.00391-17.

10. Zhang Q, Lin L, Pan Y, Chen J (2021) Characterization of Tigecycline-Heteroresistant Klebsiella pneumoniae Clinical Isolates From a Chinese Tertiary Care Teaching Hospital. Front Microbiol. 12:671153. doi: 10.3389/fmicb.2021.671153.

11. Garcia-Fulgueiras V, Magallanes C, Reyes V, Cayota C, Galiana A, Vieytes M et al (2021) In Vivo High Plasticity of Multi-Drug Resistant ST258 Klebsiella pneumoniae. Microb Drug Resist 27:1126–1130. doi: 10.1089/mdr.2020.0310.

12. Nava RG, Oliveira-Silva M, Nakamura-Silva R, Pitondo-Silva A, Vespero AC (2019) New sequence type in multidrug-resistant Klebsiella pneumoniae harboring the blaNDM-1-encoding gene in Brazil. Int J Infect Dis 79:101–103 doi: 10.1016/j.ijid.2018.11.012.

13. Meade E, Slattery MA, Garvey M (2021) Biocidal Resistance in Clinically Relevant Microbial Species: A Major Public Health Risk. Pathogens 10:598. doi: 10.3390/pathogens10050598.

14. Vats P, Kaur UJ, Rishi P (2022) Heavy metal-induced selection and proliferation of antibiotic resistance: A review. J Appl Microbiol 132:4058–4076. doi: 10.1111/jam.15492.

15. Tan Y, Zhao K, Yang S, Chen S, Li C, Han X et al (2024) Insights into antibiotic and heavy metal resistance interactions in Escherichia coli isolated from livestock manure and fertilized soil. Environ Manage 351:119935. doi: 10.1016/j.jenvman.2023.119935.

16. Mendes G, Santos ML, Ramalho JF, Duarte A, Caneiras C (2023) Virulence factors in carbapenem-resistant hypervirulent Klebsiella pneumoniae. Front Microbiol 14:1325077. doi: 10.3389/fmicb.2023.1325077.

17. Hsieh PF, Lin TL, Yang FL, Wu MC, Pan YJ, Wu SH, Wang JT (2012) Lipopolysaccharide O1 antigen contributes to the virulence in Klebsiella pneumoniae causing pyogenic liver abscess. PLoS One 7:e33155. doi: 10.1371/journal.pone.0033155.

